# Stochastic Regression and Peak Delineation with Flow Cytometry Data

**DOI:** 10.1101/2025.02.21.639492

**Authors:** Anthony J. Kearsley, Kirsten H. Parratt, Guilherme L. Pinheiro, Sandra M. Da Silva

## Abstract

Many modern molecular analysis methods utilize DNA content values as part of the measurement process, and thus, the distribution of genome copies per cell within a population of cells is important. Genome copy distributions can be measured via flow cytometry by thresholding (or “gating”) a subset of cells from which estimates of the targeted properties (e.g., genome copy number) can be calculated. This manuscript introduces a new approach that gives separate estimates of signal and noise, the former of which is used for gating and analysis, and the latter is used to quantify uncertainty. In this approach stochastic regression was used to quantify subpopulations of cells that have distinctly different genome copies per cell within a heterogenous population of *Escherichia coli* (*E. coli)* cells. By separating the signal and noise components, they can be used independently to evaluate measurement quality across different experimental conditions.

## Introduction

Molecular analytical techniques, such as quantitative polymerase chain reaction (qPCR), digital polymerase chain reaction (dPCR), and next-generation sequencing, are key contributors to microbial measurements for biotechnology, microbiome, biosurveillance, and other applications. However, to compare the data from these molecular methods with other types of microbial measurements, it is essential to establish the distribution of genome copies per cell to convert between values accurately. Flow cytometry paired with DNA-specific fluorescent dyes enables high-throughput, single-cell analysis of DNA content thereby generating plots of the distribution of DNA content across cells from a given population [1–4]. These fluorescent intensity measurements can then be converted to genome copy number estimates based on the counts associated with the peak intensities, though there are many technical considerations [5–7]. The plant research community has demonstrated standardization efforts using flow cytometry to achieve absolute quantification of genome copy number [8, 9] and benchmarks measurements against results from other methods (e.g., Feulgen microdensitometry) [10–12]. Our objective is to enable similar absolute quantifications for microbial cells.

Similar analyses can be performed for model microbial organisms (e.g., *Escherichia coli* and *Caulobacter crescentus*) with well-understood DNA replication mechanisms. Extensive and detailed biochemical descriptions of their replication can be found elsewhere, including the expected number of genome copies depending on the growth condition [13–15]. For these species with known mechanism of replication, a DNA-specific, cell-permeant fluorescent probe (e.g., Hoechst33342) can label genomes in intact cells (**Figure 1**), and the distribution of fluorescent intensities, measured by flow cytometry, is used to calculate the number of genome copies on a per-cell basis. The use of flow cytometry to evaluate genome copy distributions in different microbial cells presents several challenges. Each strain of microbe may require specific optimization [16, 17] of fluorescent probe staining in addition to other techniques to convert fluorescence intensity in genome copies. Additionally, peaks observed from non-model organisms cannot be directly correlated to genome copy numbers [18, 19]. For example, an incomplete DNA round of replication, known as multi-fork DNA replication, may result in overlapping peaks in flow cytometry histograms. When peaks overlap, the choice of the threshold value(s), referred to as “gating” in flow cytometry, can significantly influence the estimation of cells in each subpopulation as well as the distribution of genome copies. Therefore, it is critical to reduce the impact of measurement uncertainty when determining gating thresholds.

**Figure 1.**
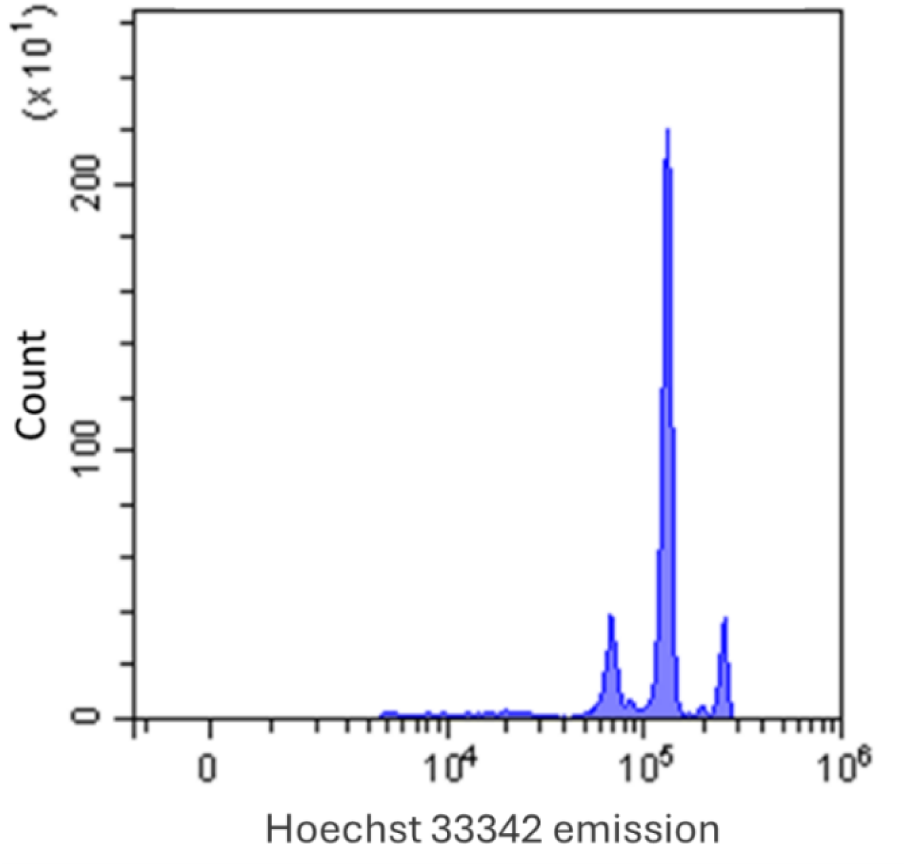
Flow cytometry histogram of *E. coli* NIST 0056 cells stained with Hoechst shows three peaks representing subpopulations of *E.coli*.

Automated gating has been repeatedly identified as important for reducing variability during flow cytometry data analysis and many methods have been proposed for different applications [20–23]. Data analysis strategies often require transforming the raw data to either histograms or an estimate of the underlying probability distribution function (PDF). Kernel density estimation (KDE) is a choice for the latter, which yields a smooth PDF estimate with a suitable kernel [24]. However, the latter doesn’t provide an inherent noise estimate. Noise in flow cytometry can arise from various sources, contributing to measurement uncertainty. It may stem from the biological sample being analyzed (e.g. true variability between cells, incomplete genome replication), the detection methods used, variations in the orientation and position of the analyte within the flow stream, or stochastic processes related to light emission and its detection. In summary, cytometers combine the inherent variability of biological populations with uncertainties introduced by the instruments[25, 26]. Applying stochastic regression to optimally constructed histograms yield estimated deterministic and stochastic components. The former can be used for analysis and gating while the latter facilitates uncertainty quantification. Many studies in the literature have mainly focused on uncertainty related to instrument characterization, such as measuring the signal-to-noise ratio under different acquisition settings. However, we hypothesized that a substantial portion of uncertainty may arise from the system’s biology processes such as the rapid genome replication in nutrient-rich media [27]. Our study demonstrates that this uncertainty can be quantified.

As a proof of concept, a test dataset was utilized to evaluate the proposed method of estimating and removing the noise contribution before enumerating events in subpopulations. *E.coli* cells were grown under defined conditions to yield a set of partially overlapping fluorescent peaks characteristic of many microbial samples. The cells were diluted to a range of experimentally relevant concentrations, the DNA was fluorescently labeled with Hoechst33342, and all other flow cytometry acquisition steps were kept constant across samples. Since all samples came from the same underlying distribution, it was hypothesized that the analysis method would give similar estimates of relative peak proportions across all samples. The objective was to evaluate the applicability of the proposed analysis method for microbial flow cytometry measurements to quantify uncertainty before estimating the distribution of subpopulations.

## Methods

*Cells: Escherichia coli* NIST0056 was cultured for 120 h at 37 °C under 120 rpm shaking at ambient atmospheric conditions. Two milliliters of the culture were spun down (10,000 x g for 2 min), followed by supernatant removal and pellet re-suspension in 2 mL phosphate-buffered saline (pH 7.2) and storage at 4 °C in preparation for analysis via flow cytometry. Details of the cell growth can be found in the supplemental material.

*Flow Cytometry Data Preprocessing*: The raw data (FCS files) were exported from the instrument and the flowAI package [28] in RStudio was used to evaluate data quality. Only data collected after the first 60 seconds were used. Background events were removed using the FlowGateNIST method [29] in Python, followed by exporting the fluorescent intensities of the remaining detected objects. Since *E. coli* is found as single cells in suspension and blank buffers with Hoechst show few objects after gating, all remaining objects were classified as “cells.” Details can be found in the supplemental material.

## Results

In **Figure 2A**, It is illustrated a flow cytometry measurement result mapped to a histogram.

**Figure 2.**
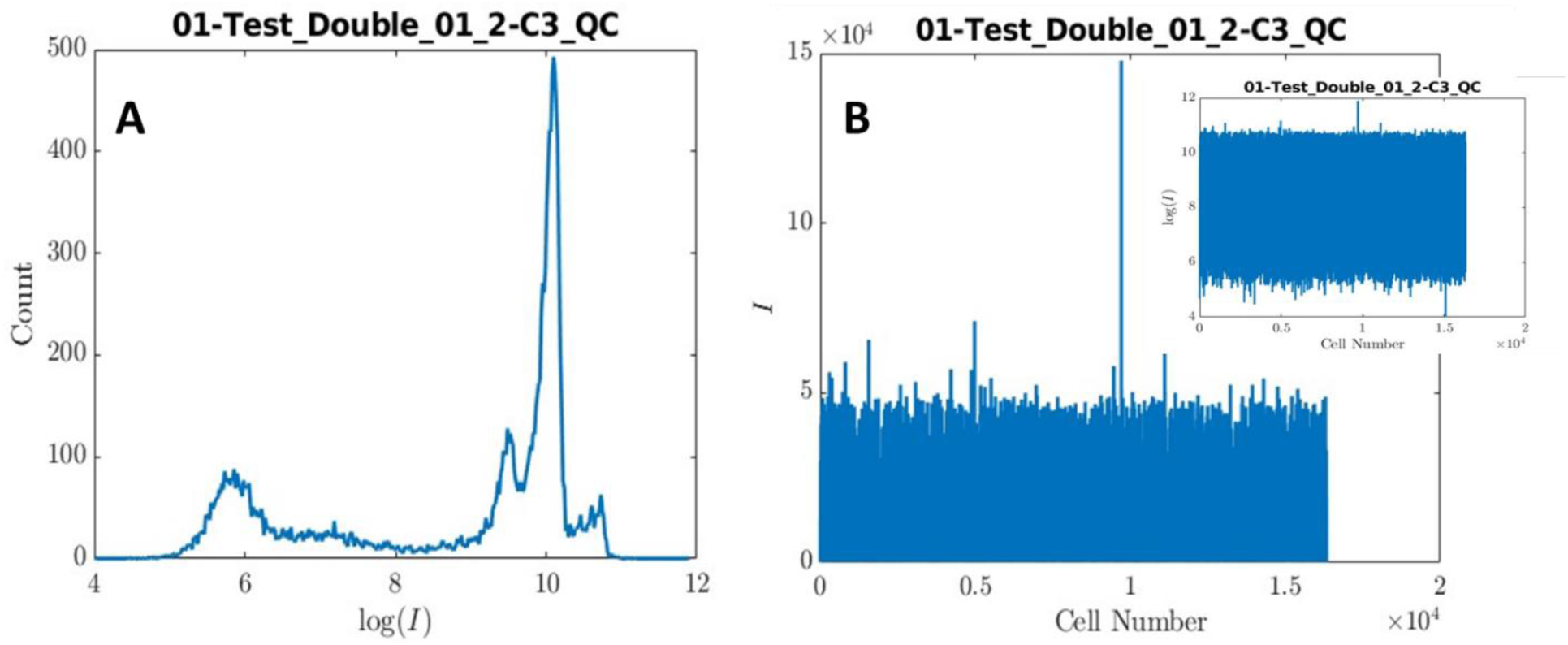
Logarithmic conversion of the data mapped to a histogram (**A**) and raw flow cytometry measurement (**B**). I, denotes the intensity associated with a given event. The figure inset in **B** has the Y-axis (I) converted to a logarithmic scale.

As a representative example of the first step of the data analysis, the histogram shown in **Figure 1A** was constructed with the raw data described in **Figure 2B**. The vertical axis of **Figure 2A** describes cell number while the horizontal denotes fluorescence intensity (*I*) corresponding to the fluorescence from the DNA-binding dye. Intensities range from 0 to approximately 15 × 10^4^ as shown in **Figure 2B**. To appropriately scale the data we apply log_e_(·) (henceforth denoted as log(·)) to each of the intensities. Applying a logarithmic scaling to the data in **Figure 2B** results in the data shown in the inserted figure **(Figure 2B)**. At this point, the data are suitable to be converted to the histogram described in **Figure 2A**, which shows the cell count as a function of log(*I*), and to detect biologically significant features.

To this end, the vertical axis in **Figure 2B** must be *binned* by partitioning it into *N* subintervals. A histogram is then constructed by counting the number of cells that fall within each sub-interval and graphically represented by letting the mid-point of each sub-interval be the horizontal coordinate and the number of cells within that sub-interval be the vertical coordinate. For example, if there are *j* cells in the sub-interval [*y_n_*_−1_*, y_n_*], then the corresponding point on the histogram is given in **Equation 1**. Histograms with *N* = 20 and *N* = 2000 are shown in **Figure 3**, where it is seen that the number of bins affects histogram quality.

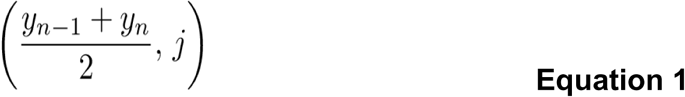

**Figure 3.**
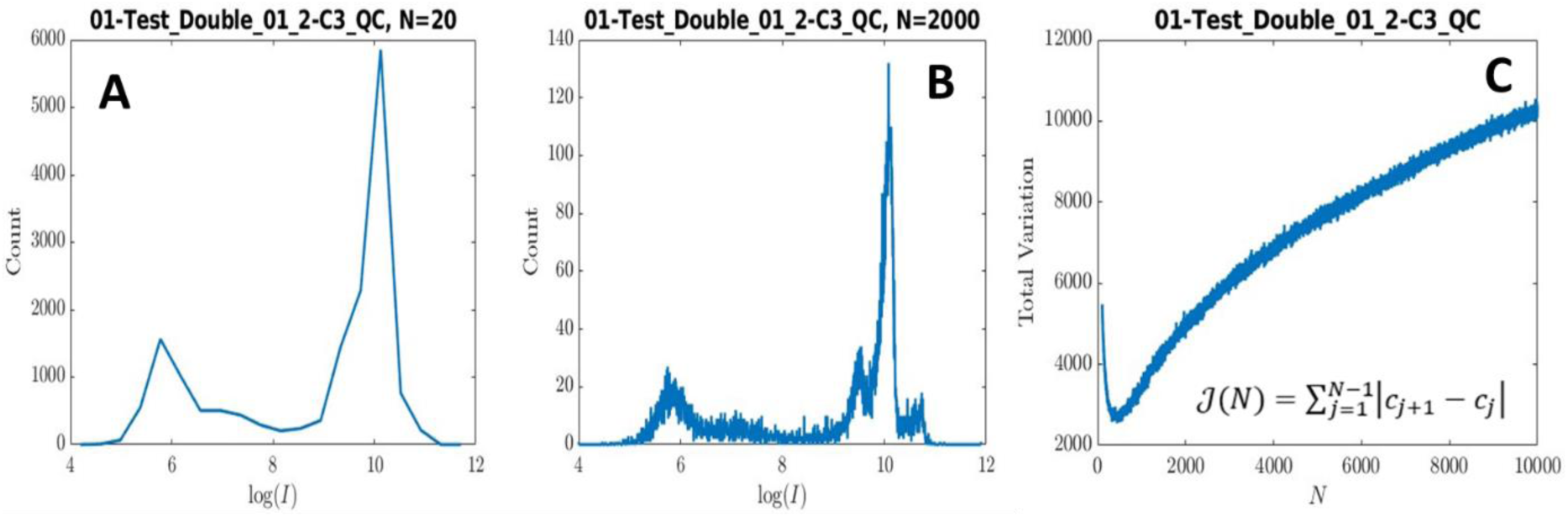
Histogram of the raw flow cytometry measurement (Figure 2A) is presented in two histogram resolutions: N = 20 (**A**) and N = 2000 (**B**). In addition, graph **C** represents the data depicted in **A** and **B.** The **Equation 2** generated the curve.

Key features will be missed if there are too few bins as demonstrated in **Figure 3A**. For instance, smaller peaks directly prior to and posterior to the largest peak around log(*I*) = 10 are no longer apparent. On the other hand, using too many bins may capture spurious fluctuations, as shown in **Figure 3B**. Furthermore, the number of bins greatly affects the count associated with each log(*I*) in the corresponding histograms. In **Figure 3A** counts/bin range from 0 to 6000, while on **Figure 3B**, counts/bin range from 0 to 140. Therefore, to produce accurate histograms it is crucial to optimally choose the number of bins *N*. Since the goal is to generate histograms as smoothly as possible, we chose *N* such that the total variation is at a minimum.

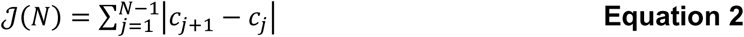

The *c_j_* in **Equation 2** denotes the number of cells associated with log(*I_j_*). A graph of the objective function is shown in **Figure 2C**. This is a bounded function of one discrete variable over a bounded interval, so a global minimum probably exists, and performing a direct search shows that, in this case, it attains a minimum at *N* = 440.

Each of the histograms in this work has its own optimal value of *N*, ranging from 440 to 948. The histogram described in **Figure 2A** has *N* = 440 bins. This is the histogram corresponding to the value of *N* that minimizes the total variation and maximizes smoothness to efficiently estimate peak boundaries. To separate *signal, i*.*e*., the deterministic component of the histogram, from noise, we apply stochastic regression. Due to the noise inherent in the instrument, it is not a surprise to see some noise even after choosing *N* to minimize the total variation (N = 440).

The stochastic regression approach to the problem was developed by Kearsley, Gadhyan, and Wallace [30] to separate signal from noise in matrix-assisted laser desorption ionization reflectron time-of-flight (MALDIRTOF) mass spectrometry data and was used to separate signal from noise in biological field effect transistor (Bio-FET) data [31]. Melara *et al.* optimized this algorithm for Bio-FET measurements [32]. Stochastic regression centers around modeling the histogram with a stochastic differential equation (SDE):

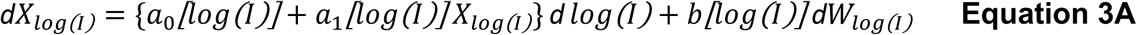

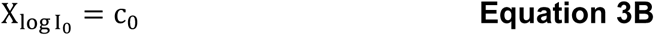

The first term {a_0_[log(I)] + a_1_[log(I)] X_log(I)_} d log(I) models the deterministic component of the measurement, *i*.*e*., the approximate signal. The second term *b*[log(*I*)] d*W*_log(*I*)_ models the stochastic component, representing the noise. The terms of the former are called the *drift* terms, and the term of the latter is called the diffusion term; the stochastic drift models the change of the average value of a stochastic process, while the diffusion models density fluctuations. Taken together models of the form (**Equation 3A**) are known as *drift-diffusion* models and are employed in a variety of applications. The drift coefficients *a*_0_[log(*I*)] and *a*_1_[log(*I*)] were estimated using local weighted regression, and the diffusion coefficient *b*[log(*I*)] was computed using maximum likelihood estimation [30]. We note that calculating the drift and diffusion coefficients depends on the size of the *averaging window*. If this window is either too large or too small, it leads to a poor approximation of the signal. For this work, the averaging window was chosen in an optimal way in accordance with Melara, *et al* [32].

The approximate signal was obtained by replacing the coefficients in **Equation 3a** with those found through local weighted regression and maximum likelihood estimation and then applying Euler’s method after omitting the diffusion coefficient. Noise was calculated by discretizing the Brownian motion term, and multiplying the result by the coefficient *b*[log(*I_j_*)] for each index *j* except 0 (since we are taking our initial condition to be exact). The results are shown in **Figure *4***. The stochastic regression does an excellent job separating signal from noise, which aids analysis in primarily two ways: 1-The data are now in a suitable form for peak delineation and 2) The noise estimate allows us to estimate the uncertainty in the number of cells associated with each peak, along with the relative percentage of cells associated with each peak.

**Figure 4.**
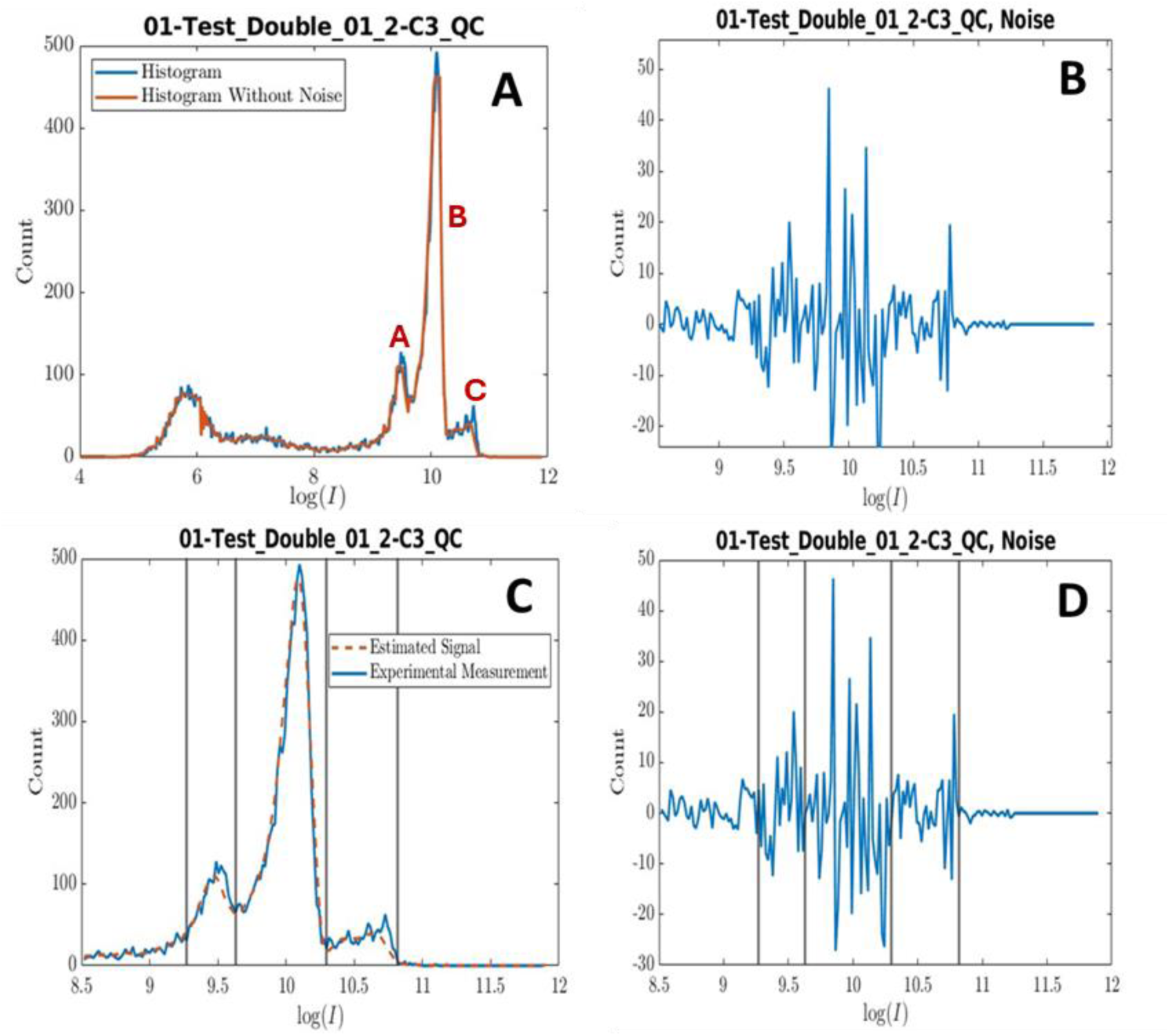
The resulting histogram after applying stochastic regression (orange) overlapped with the original histogram (blue) **(A)**, experimental measurement noise (**B**). Estimated signal after peak delineation (orange), experimental measurement (blue) **(C),** and corresponding noise location (**D**).

**Figure 5.**
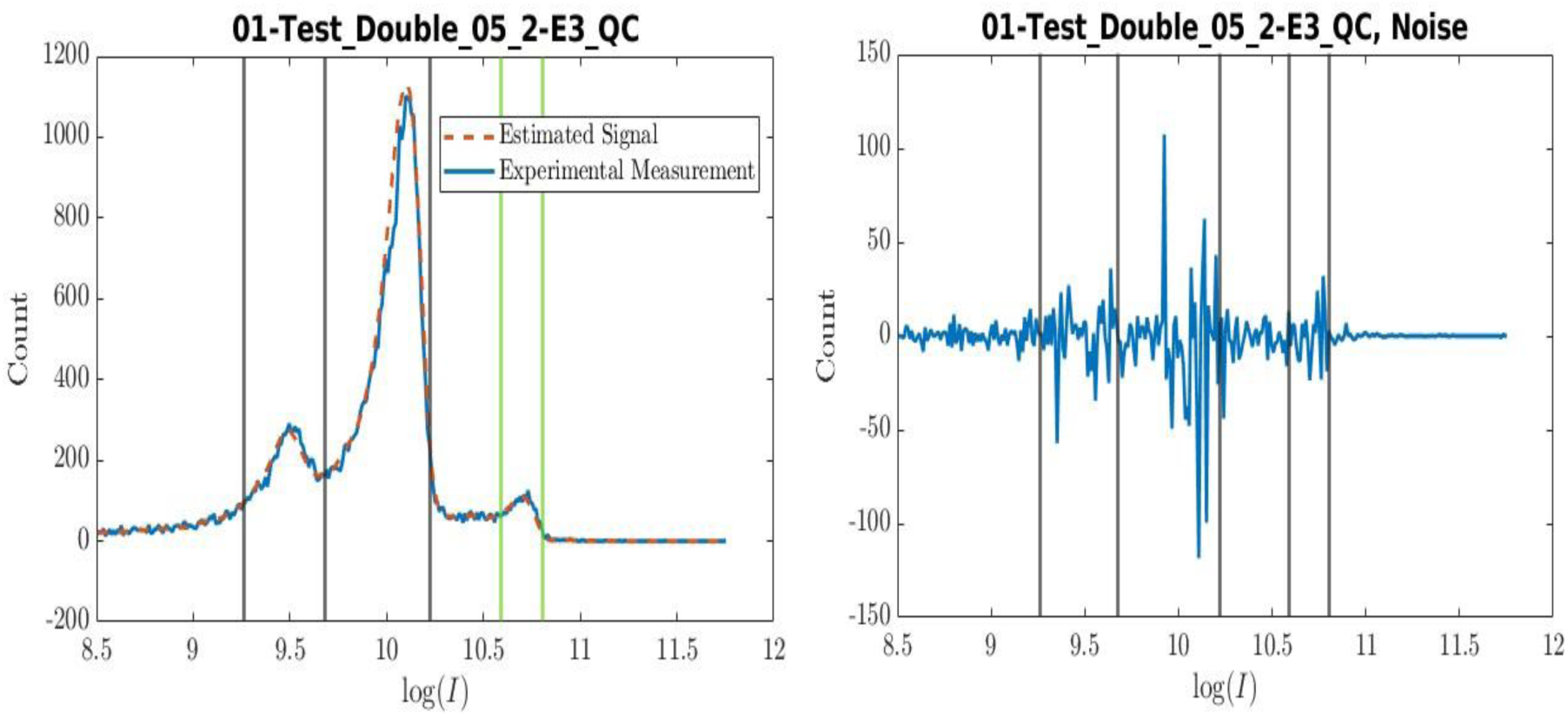
Estimated signal after peak delineation (orange), experimental measurement (blue) (A), and corresponding noise location (**B**) to test the robustness of the stochastic regression and SavitzkyGolay approaches.

Indeed, this is a very strong merit of the method since accurate quantitative analysis requires uncertainty quantification, and noise estimates are typically unavailable when solely traditional smoothing techniques are applied. Although stochastic regression gives estimates of signal and noise, it is limited by the noise in the data and error inherent in discretizing **Equation 3**. To make the data even more amenable to our peak delineation techniques, we applied a Savitzky-Golay filter [35] with a small window to improve the efficacy of the peak delineation.

The algorithm is based on the intuitive observation that peaks are separated by the bottom of valleys. On **Error! Reference source not found.4A**, there are three peaks of interest: one around log(*I*) = 9.5 (“peak A”), a second around log(*I*) = 10 (“peak B”), and a third around log(*I*) = 10.6 (“peak C”). The peak around log(*I*) = 5.75 corresponds to a non-biologically significant feature (background events with low scatter intensities, purposefully collected on the instrument to confirm that all cell events were above the acquisition threshold); hence, in the proceeding analysis, everything below log(*I*) = 8.5 will be truncated. The natural boundary between peaks A and B is the trough minimum. Mathematically this can be described as a point at which convexity is at a local maximum. To delineate peak A and B, we draw their boundaries at points of maximum convexity between peaks. A similar strategy also applies to finding where peak A begins and peak C ends. A natural location of the end of peak C is at the “elbow” around log(*I*) = 11. This is also a point at which convexity reaches a local maximum, consequently, to find where peak A begins and the peak C ends, we find points of maximum convexity over bounded intervals that are highly probable to contain the peak beginnings and endings. For example, when searching for the end of peak C we maximized convexity over the interval 10.7 and 11.5 **(Figure 4A)**. If we had an infinitely smooth amount of data, we could locate the local minimums within machine precision; however, this is not the case. Since we used a finite difference approximation to estimate curvature at points of interest, locations of maximum convexity are only approximate and subject to both noise and discretization error.

Our algorithm does an excellent job at delineating peaks; the boundaries agree with where one would expect based on visual inspection (**Error! Reference source not found.4C)**. Stochastic regression and application of Savitzky-Golay played a central role in smoothing the data, so that accurate curvature estimates could be obtained with finite differences, thereby improving the accuracy with which peak boundaries were detected. This is true especially of peak C. Notice that in the experimental measurement from approximately log(*I*) = 10.3 to log(*I*) = 10.8 it is difficult to distinguish between parts that correspond to noise and parts that are biologically significant. By applying stochastic regression, we can separate the estimated stochastic component from estimated deterministic component. Thus, we conclude that the three spikes between log(*I*) = 10.3 and log(*I*) = 10.8 in the experimental measurement are random fluctuations due to noise, rather than biologically significant features. Note that the estimated deterministic component was used to calculate the peak boundaries (as for each of the measurements herein). In the orange curve there is an unambiguous peak that begins at log(*I*) = 10.29 and ends at log(*I*) = 10.81.

To illustrate the robust nature of our algorithm, we examined the measurement shown in **Error! Reference source not found.**. Visual inspection reveals that this histogram is qualitatively different than the one examined previously, although peak C ends around log(*I*) = 10.80, which is very close to the end of peak C in the previous case, this time it begins around log(*I*) = 10.58. This can be accounted for by searching for a maximum of convexity between log(*I*) = 10.46 and log(*I*) = 10.68, which yields a local maximum of convexity around log(*I*) = 10.58. Thus, there are five relevant points at which convexity reaches a local maximum, rather than four in the previous case. Peak A is between the first two (9.26 ≤ log(*I*) ≤ 9.68), peak B is between the second and the third (9.68 ≤ log(*I*) ≤ 10.22), and peak C is between the *fourth* and *fifth* highlighted in green (10.58 ≤ log(*I*) ≤ 10.80). Before concluding this example, we would like to comment on the location of the third local convexity maximum around log(*I*) = 10.22. Visual inspection may suggest that this peak boundary should have been placed slightly to the right around log(*I*) = 10.30. The nature of the curve from log(*I*) = 10.09 to approximately log(*I*) = 10.30 is such that its curvature is particularly pronounced along this section. This, along with the discrete nature of our data and traces of noise, renders the problem of finding the point along this section of the curve at which convexity attains a local maximum ill-conditioned. Since the problem of locating the point along this section of the curve at which convexity reaches a local maximum is ill-conditioned, it is expected that our algorithm’s location of this point may not agree exactly with our intuition.

These two examples show that we can locate peak boundaries using curvature information by applying stochastic regression and SavitzkyGolay. Finally, these peak boundaries can be used to calculate the *total* number of cells associated with *all the peaks* (N_i_) and determine the relative percent of cells associated with each of them **(Equation 4**).

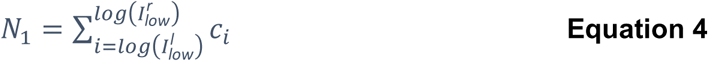

Where N_1_ is the total number of cells associated with the first peak, log(*I*_low_^l^) denotes the left boundary of peak A, log(*I*_low_^r^) denotes the right boundary of peak A, and *c_j_* denotes the count corresponding to intensity log(*I_j_*). Then if there are *N*_2_ and *N*_3_ cells that are similarly associated with the relative percent associated with each of the peaks can be defined as

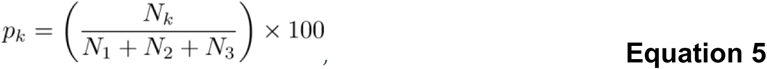

for *k* = 1, 2, 3. The relative percentages associated with each peak are given in **Table *1*** and graphed in **Figure *6***, and the mean and standard deviation of relative percentages are given in **Table *2***. The amount of noise associated with each of the peaks is also of interest, so we have calculated the noise associated with each peak by taking the *l*_1_ norm of the estimated stochastic component of each histogram.

**Figure 6.**
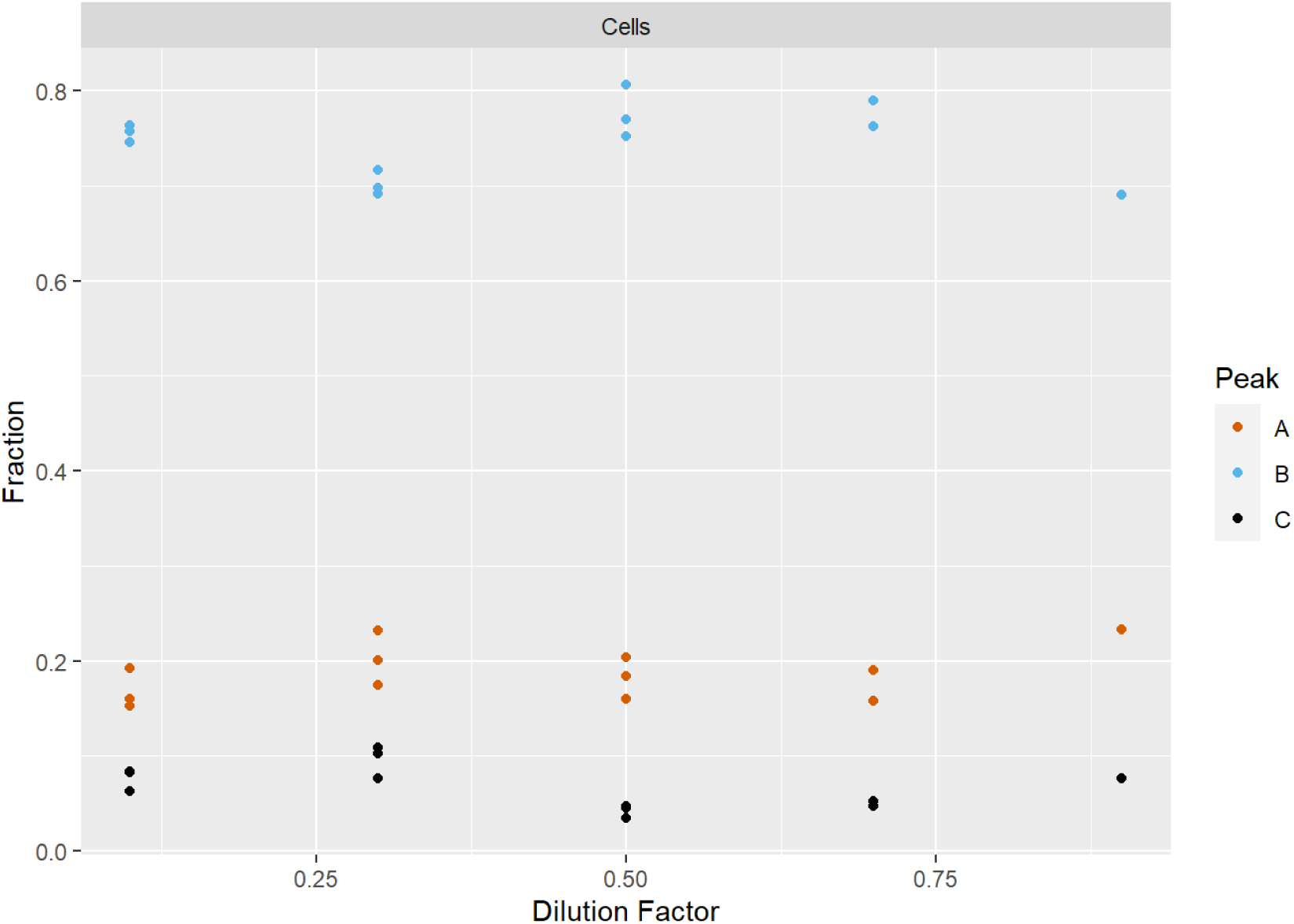
The fraction of objects assigned to each peak is not dependent on the dilution factor.

**Table 1.** Peak statistics. Columns 1 to 3 are the count associated with each peak, column 4 represents the total count associated with each histogram, and columns 5 to 7 represent the relative percent associated with each peak. The numbers in this table were calculated using the estimated deterministic component of each histogram.

| Measurement | Low Count | Medium Count | High Count | Total Count | Low % | Medium % | High % |
| --- | --- | --- | --- | --- | --- | --- | --- |
|  | (1) | (2) | (3) | (4) | (5) | (6) | (7) |
| 01-Control Hoechst Ctrl 1-A2 QC | 11 075.29 | 42 056.63 | 4767.83 | 57 899.74 | 19.13 | 72.64 | 8.23 |
| 01-Test Double 01 1-C1 QC | 1908.36 | 7405.09 | 618.97 | 9 932.42 | 19.21 | 75.55 | 6.23 |
| 01-Test Double 01 2-C3 QC | 1669.08 | 7899.10 | 861.77 | 10 429.95 | 16.00 | 75.73 | 8.26 |
| 01-Test Double 01 3-C5 QC | 1484.45 | 7438.87 | 861.04 | 9739.36 | 15.24 | 76.38 | 8.38 |
| 01-Test Double 03 1-D1 QC | 5972.50 | 20 831.91 | 3052.96 | 29 857.37 | 20.00 | 69.77 | 10.23 |
| 01-Test Double 03 2-D3 QC | 5730.80 | 23 574.36 | 3587.92 | 32 893.08 | 17.42 | 71.67 | 10.91 |
| 01-Test Double 03 3-D5 QC | 5801.13 | 17 265.85 | 1906.54 | 24 973.51 | 23.23 | 69.14 | 7.63 |
| 01-Test Double 05 1-E1 QC | 7470.10 | 31 317.35 | 5008.52 | 43 795.88 | 17.06 | 71.51 | 11.44 |
| 01-Test Double 05 2-E3 QC | 8027.24 | 29 688.9 | 4495.32 | 42 211.46 | 19.02 | 70.33 | 10.65 |
| 01-Test Double 05 3-E5 QC | 8086.01 | 40 673.17 | 5898.15 | 54 657.32 | 14.79 | 74.41 | 10.79 |
| 01-Test Double 07 1-F1 QC | 10 528.71 | 52 562.97 | 8699.04 | 71 790.72 | 14.67 | 71.09 | 11.15 |
| 01-Test Double 07 2-F3 QC | 11 474.45 | 45 952.94 | 7208.66 | 64 626.04 | 17.75 | 71.09 | 11.15 |
| 01-Test Double 09 1-G1 QC | 20 374.32 | 60 373.18 | 9537.33 | 90 284.84 | 22.57 | 66.87 | 10.56 |

**Table 2.** Mean (µ) and standard deviation (σ) of count percentages associated with the low, medium, and high intensity peaks depicted in Table 1.

|  | Low % | Medium % | High % |
| --- | --- | --- | --- |
| $\mu$ | 18.70 | 74.38 | 6.92 |
| $\sigma$ | 2.63 | 3.69 | 2.33 |

Mathematically the *l*_1_ norm of a vector (*w*_1_*,…, w_n_*)*^T^* is given by

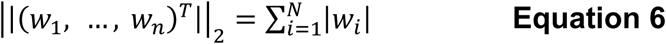

The noise associated with each peak has been tabulated in **Table *3***. To evaluate the relative impact of removing this noise, the noise count per peak was divided by the sum of the noise count and cell count per peak (**Figure *7***).

**Figure 7.**
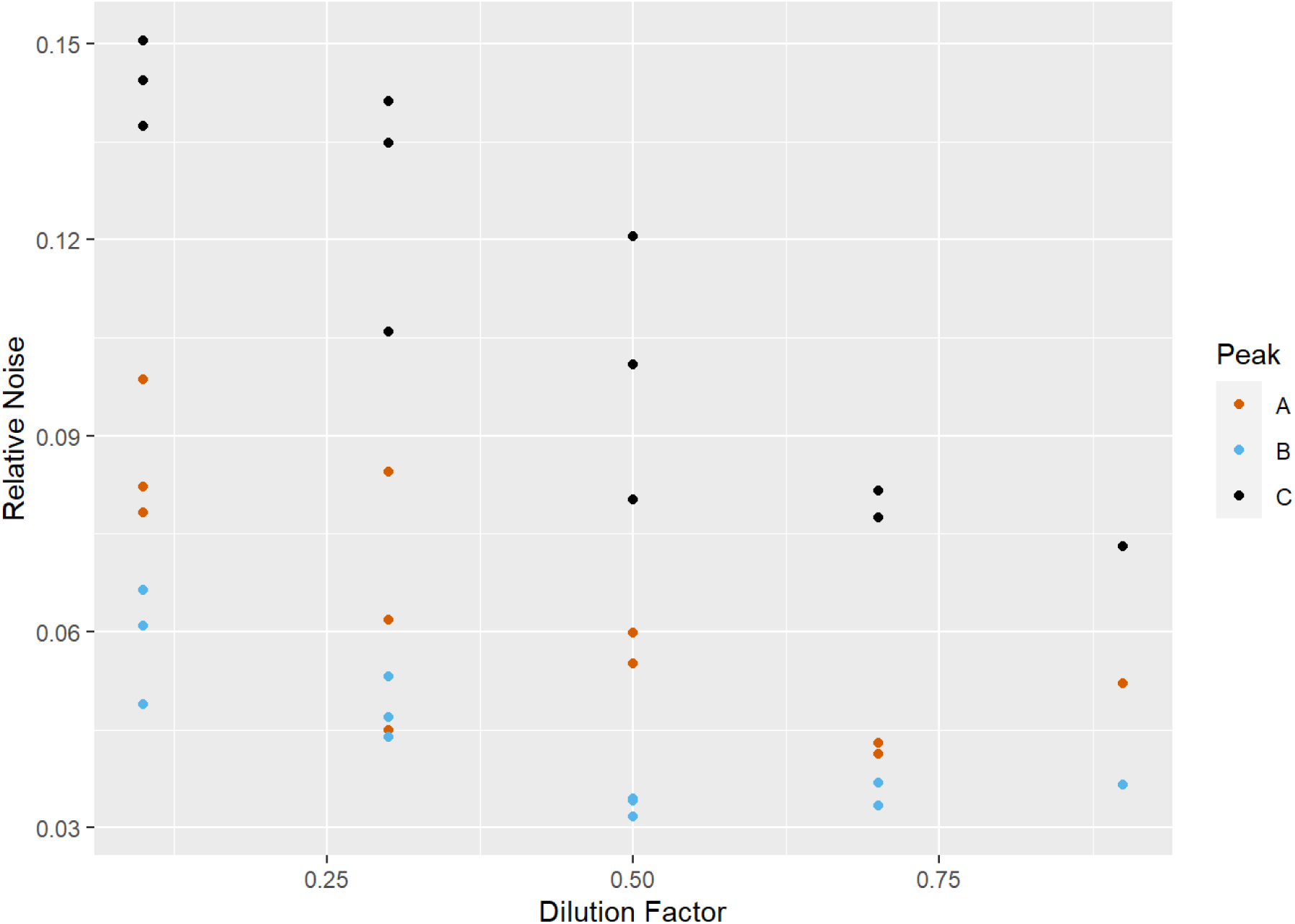
The relative fraction of events in a peak that were classified as noise by our regression analysis. Dilution factor = 1 corresponds to no dilution was performed whereas dilution factor = 0.1 corresponds to a 10-fold dilution.

**Table 3.** The calculated noise associated with each of the peaks is given by the l_1_ norm of the stochastic component of each peak signal.

| Measurement | Low Count | Medium Count | High Count | Total Count |
| --- | --- | --- | --- | --- |
| 01-Control Hoechst Ctrl 1-A2 QC | 606.69 | 1309.87 | 543.07 | 2459.86 |
| 01-Test Double 01 1-C1 QC | 208.84 | 526.44 | 109.60 | 844.89 |
| 01-Test Double 01 2-C3 QC | 141.73 | 405.83 | 137.22 | 684.78 |
| 01-Test Double 01 3-C5 QC | 133.03 | 481.85 | 137.74 | 752.62 |
| 01-Test Double 03 1-D1 QC | 281.35 | 1026.16 | 475.95 | 1783.47 |
| 01-Test Double 03 2-D3 QC | 3777.08 | 1324.14 | 425.01 | 2126.86 |
| 01-Test Double 03 3-D5 QC | 534.79 | 793.05 | 313.53 | 1641.38 |
| 01-Test Double 05 1-E1 QC | 264.76 | 1115.96 | 165.87 | 1546.65 |
| 01-Test Double 05 2-E3 QC | 467.80 | 1047.67 | 240.90 | 1756.37 |
| 01 Test Double 05 3-E5 QC | 514.69 | 1329.55 | 192.68 | 2036.92 |
| 01 Test Double 07 1-F1 QC | 472.21 | 1818.51 | 309.15 | 2599.78 |
| 01 Test Double 07 2-F3 QC | 493.65 | 1758.27 | 238.06 | 2489.99 |
| 01 Test Double 09 1-G1 QC | 1119.74 | 2289.16 | 526.27 | 3935.17 |

Due to the nature of the experiments, variation in the counts associated with each of the peaks is expected. However, the relative percentage data should be consistent across samples. **Table *2*** gives the mean and standard deviation of the relative percent associated with each peak. The latter gives a measure of how much we can expect the relative percentages to deviate from their mean values. For the data presented herein, all relative percentages are within two standard deviations of their mean value.

## Discussion

This study presents an innovative method for analyzing *E. coli* subpopulations, which may be applicable to other microbial cells, by providing distinct estimates for both signal and noise. This approach aims to quantify uncertainty and better understand the distribution of these subpopulations. The primary goal is to assess the feasibility of this analytical technique in microbial flow cytometry measurements, specifically focusing on quantifying the noise prior to estimating the distribution of the subpopulations.

Our approach was tested using a dataset that reflects the typical characteristics of microbial flow cytometry results. Since all the test samples were drawn from the same underlying population, the deterministic estimates of the relative percentages of subpopulations were found to be consistent across all samples (**Figure *6***). The estimates’ variation was consistent across all three peaks (**Table 2**). Although the gates chosen here exclude a small number of events at either end of the histogram, these excluded events represent only a very small proportion of the total number. In contrast to other methods, our method also provides an uncertainty estimate based on the stochastic component of the genome copy distributions. Furthermore, gating methods are applied after mapping raw data to either a histogram or PDF. The latter is typically done using KDE, and although the resulting PDF is smooth with an appropriately chosen kernel, this does not provide a noise estimate. It has been shown that applying stochastic regression to optimally constructed histograms separate signal from noise, which facilitates quantitative analysis and gives a way to quantifying uncertainty. This uncertainty estimate was proportionally larger for peaks with smaller counts, an expected relationship especially if peaks are isolated. A deeper understanding of this aspect of measurement uncertainty is valuable when comparing genome copy estimates across different cytometers or even other measurement technologies such as dPCR, a technique that, in principle, can be used in conjunction with flow cytometry to assign genome copies. The mathematical techniques demonstrated here could also be extended to flow cytometry measurements other than genome copy.

Many automated gating methods have been applied to flow cytometry data with great success at reducing variability from operator-specific analysis biases. These methods appear well suited to analyze flow cytometry data after uncertainty quantification, similar to how peak delineation was performed here by applying stochastic regression and Savitzky Golay. In these experiments and in most cytometry, contributions to noise are inherently probabilistic and thus a stochastic model seems more appropriate than a deterministic model that does not incorporate probability. In the literature, there many papers that compare gating strategies to assess relative performance but there generally is not a “ground truth” obtained from an orthogonal measurement that can be compared (ground truths might instead relate to data quality issues, manual gating results, or clinical status) [33, 34]. NIST is currently developing whole microbial cell reference materials characterized for cell object count and genome copy count which can be compared between methods and thus begin to evaluate which gating methods arrive at results closer to ground-truth values.

## Conclusion

Genome copy measurements are important for several burgeoning applications in microbiology, but there are challenges in obtaining accurate results. The proposed method offers a new way to evaluate flow cytometry measurements of genome copy distributions and will be especially critical and important for samples where less distinct peaks are observed. In summary, cell DNA patterns are complex in microbial cells as they depend on growth conditions. The ability to quantify genome copies and associated uncertainties play an important role in the accuracy of DNA-based measurements underpinning significant advances in biotechnology.

## Supporting information

Sup. Mat.

## Acknowledgements

The authors thank Ryan Evans (NIST, Applied and Computational Mathematics Division) for substantial discussions and data analysis.

## Disclaimer

Certain commercial equipment, instruments, or materials are identified in this presentation to foster understanding. Such identification does not imply recommendation or endorsement by the National Institute of Standards and Technology, nor does it imply that the materials or equipment identified are necessarily the best available for the purpose.

## Conflict of Interest

The authors declare that they have no conflict of interest.

## Notes

https://doi.org/10.18434/mds2-3119

### Competing Interest Statement

The authors have declared no competing interest.

## References

1. Akerlund, T., K. Nordstrom, and R. Bernander, Analysis of cell size and DNA content in exponentially growing and stationary-phase batch cultures of Escherichia coli. J Bacteriol, 1995. 177(23): p. 6791–7.

2. Muller, S., Modes of cytometric bacterial DNA pattern: a tool for pursuing growth. Cell Prolif, 2007. 40(5): p. 621–39.

3. Allman, R., et al., Characterization of bacteria by multiparameter flow cytometry. J Appl Bacteriol, 1992. 73(5): p. 438–44.

4. Allman, R., T. Schjerven, and E. Boye, Cell cycle parameters of Escherichia coli K-12. J Bacteriol, 1991. 173(24): p. 7970–4.

5. Dolezel, J. and J. Greilhuber, Nuclear genome size: are we getting closer? Cytometry A, 2010. 77(7): p. 635–42.

6. Dolezel, J., S. Sgorbati, and S. Lucretti, Comparison of 3 DNA Fluorochromes for Flow Cytometric Estimation of Nuclear-DNA Content in Plants. Physiologia Plantarum, 1992. 85(4): p. 625–631.

7. Suda, J. and I.J. Leitch, The quest for suitable reference standards in genome size research. Cytometry A, 2010. 77(8): p. 717–20.

8. Koutecky, P., et al., Best practices for instrument settings and raw data analysis in plant flow cytometry. Cytometry Part A, 2023.

9. Sliwinska, E., et al., Application-based guidelines for best practices in plant flow cytometry. Cytometry Part A, 2022. 101(9): p. 749–781.

10. Temsch, E.M., et al., Reference standards for flow cytometric estimation of absolute nuclear DNA content in plants. Cytometry Part A, 2022. 101(9): p. 710–724.

11. Galbraith, D.W., Validation of crowd-sourced plant genome size measurements. Cytometry Part A, 2022. 101(9): p. 703–706.

12. Baranyi, M. and J. Greilhuber, Flow cytometric and Feulgen densitometric: Analysis of genome size variation in Pisum. Theoretical and Applied Genetics, 1996. 92(3-4): p. 297–307.

13. McAdams, H.H. and L. Shapiro, A bacterial cell-cycle regulatory network operating in time and space. Science, 2003. 301(5641): p. 1874-7.

14. Gitai, Z., M. Thanbichler, and L. Shapiro, The choreographed dynamics of bacterial chromosomes. Trends Microbiol, 2005. 13(5): p. 221–8.

15. Thanbichler, M., P.H. Viollier, and L. Shapiro, The structure and function of the bacterial chromosome. Curr Opin Genet Dev, 2005. 15(2): p. 153–62.

16. Davey, H. and S. Guyot, Estimation of Microbial Viability Using Flow Cytometry. Curr Protoc Cytom, 2020. 93(1): p. e72.

17. Müller, S. and G. Nebe-von-Caron, Functional single-cell analyses: flow cytometry and cell sorting of microbial populations and communities. Fems Microbiology Reviews, 2010. 34(4): p. 554–587.

18. Bernander, R., T. Stokke, and E. Boye, Flow cytometry of bacterial cells: comparison between different flow cytometers and different DNA stains. Cytometry, 1998. 31(1): p. 29–36.

19. Lieder, S., et al., Subpopulation-proteomics reveal growth rate, but not cell cycling, as a major impact on protein composition in KT2440. Amb Express, 2014. 4.

20. Finak, G., et al., OpenCyto: An Open Source Infrastructure for Scalable, Robust, Reproducible, and Automated, End-to-End Flow Cytometry Data Analysis. Plos Computational Biology, 2014. 10(8).

21. Mair, F., et al., The end of gating? An introduction to automated analysis of high dimensional cytometry data. Eur J Immunol, 2016. 46(1): p. 34–43.

22. Smith, T.W., P. Kron, and S.L. Martin, flowPloidy: An R package for genome size and ploidy assessment of flow cytometry data. Applications in Plant Sciences, 2018. 6(7).

23. Koch, C., et al., Cytometric fingerprints: evaluation of new tools for analyzing microbial community dynamics. Frontiers in Microbiology, 2014. 5.

24. Lugli, E., M. Roederer, and A. Cossarizza, Data analysis in flow cytometry: the future just started. Cytometry A, 2010. 77(7): p. 705–13.

25. Arriaga, E.A., Determining biological noise via single cell analysis. Anal Bioanal Chem, 2009. 393(1): p. 73–80.

26. Wood, J.C., Fundamental flow cytometer properties governing sensitivity and resolution. Cytometry, 1998. 33(2): p. 260–6.

27. Giesecke, C., et al., Determination of background, signal-to-noise, and dynamic range of a flow cytometer: A novel practical method for instrument characterization and standardization. Cytometry A, 2017. 91(11): p. 1104–1114.

28. Monaco, G., et al., flowAI: automatic and interactive anomaly discerning tools for flow cytometry data. Bioinformatics, 2016. 32(16): p. 2473–80.

29. Ross, D., Automated analysis of bacterial flow cytometry data with FlowGateNIST. PLoS One, 2021. 16(8): p. e0250753.

30. Kearsley, A.J., Y. Gadhyan, and W.E. Wallace, Stochastic regression modeling of chemical spectra. Chemometrics and Intelligent Laboratory Systems, 2014. 139: p. 26–32.

31. Evans, R.M., A. Balijepalli, and A.J. Kearsley, Transport Phenomena in Biological Field Effect Transistors. Siam Journal on Applied Mathematics, 2020. 80(6): p. 2586–2607.

32. Melara, L.A., Evans, R. M., Cho, S., Balijepalli, A., Kearsley, A. J., Optimal Bandwidth Selection in Stochastic Regression of Bio-FET Measurements. submitted.

33. Aghaeepour, N., et al., *Critical assessment of automated flow cytometry data analysis techniques (vol 10, pg 228*, *2013)*. Nature Methods, 2013. 10(5): p. 445–445.

34. Emmaneel, A., et al., PeacoQC: Peak-based selection of high quality cytometry data. Cytometry Part A, 2022. 101(4): p. 325–338.

35. Savitzky, A.; Golay, M.J.E. (1964). “Smoothing and Differentiation of Data by Simplified Least Squares Procedures”. Analytical Chemistry. 36 (8): p. 1627–1639.

