## Supplementary material for "Stochastic Regression and Peak Delineation with Flow Cytometry Data": Sup. Mat.

Data availability statement: Sample data supporting the findings of this study are available at https://doi.org/10.18434/mds2-3119

**Supplemental Material**

*Cells:* One Biobank bead (Pro-lab Diagnostics - cat# P.170) containing cells was removed from -80 $^{o}$C and placed into a 15 mL conical tube containing 5 mL tryptic soy broth (TSB, BD Bacto #211825t). The tube was incubated overnight (~16 h) at 37 $^{o}$C under 180 rpm shaking at ambient atmosphere. One milliliter of the overnight culture was spun down (10,000 x g) for 2 min, supernatant was removed, and the pellet was re-suspended in 1 mL sterile water. Cells were diluted to OD600 = 0.5 (≈ 4 x $10^8$ cells/mL) and further diluted 100-fold to prepare the inoculum. Then, 62.5 µL of the inoculum (≈ 250,000 cells) was added to 5 mL TSB to obtain ≈ 50,000 cells per mL.

The optical density at 600 nm ($OD_{600}$) of the suspension was measured in phosphate buffered saline (PBS, pH 7.4). The sample was diluted to generate five concentration levels ("dilution factors") spanning one order of magnitude in the test dataset. Suspensions were stained with 4 µmol/L Hoechst33342 (DNA stain, ThermoFisher, cat#62249) and a 1:5000 dilution of CellBriteFix640 (Cell body stain, Biotium, cat30089) stock solution prepared per manufacturer's protocol, and incubated at 37 °C for 30 min. Stained samples were diluted 1:10 into PBS then analyzed on the flow cytometer (CytoFLEX LX, Beckman Coulter) at a rate of 30 µL/minute for 2 minutes per sample with the following cytometer acquisition conditions: FSC = 100, SSC-H Threshold = 5,000, NUV450 = 50 (acronyms correspond to channel names provided by the manufacturer). Thus, the same volume was acquired for all samples with purposefully different numbers of cells measured. Three replicates were collected per cell concentration. Two files were excluded because clogs during file acquisition resulted in minimal events detected.

*Flow Cytometry Data Preprocessing*: Raw data (fcs files) were exported off the instrument and preprocessed with RStudio. Quality control (QC) was run according to *flowAI package* with a flow_auto_qc function [25], and only data from analyses performed after the first 60 seconds of each data file were exported for further analysis. Files were then opened in Python, and “background” events were removed following the workflow described in FlowGateNIST [26]. For each data file, the fluorescent intensities of all remaining detected objects were exported. Since this *E. coli* is known to be single cells in suspension (microscopy, not shown) and blank buffer with Hoechst shows very few objects after gating, all objects are deemed “cells”.
